# Genome-resolved meta-omics unveils rock-hosted lifestyle of enigmatic DPANN archaea

**DOI:** 10.1101/2023.06.16.545063

**Authors:** Hinako Takamiya, Mariko Kouduka, Shingo Kato, Hiroki Suga, Masaki Oura, Tadashi Yokoyama, Michio Suzuki, Masaru Mori, Akio Kanai, Yohey Suzuki

## Abstract

Recent successes in the cultivation of DPANN archaea with their hosts have demonstrated an episymbiotic lifestyle, whereas the lifestyle of DPANN archaea in natural habitats remains controversial. A free-living lifestyle is speculated in oxygen-deprived fluids circulated through rock fractures, where apparent hosts of DPANN archaea are lacking. Alternatively, DPANN archaea may be isolated from their hosts attached to rock surfaces. To understand the ecology of rock-hosted DPANN archaea, rocks rather than fluids should be directly characterized. Here, we show the dominance of Pacearchaeota, one of the widespread and enigmatic lineages of DPANN archaea, in a deep-sea hydrothermal vent chimney. Metagenomic analysis of the rock sample revealed a symbiotic lifestyle of the chimney Pacearchaeota, based on the lack of biosynthetic genes for nucleotides, amino acids, cofactors, and lipids. Genome-resolved metaproteomic analysis clarified the co-occurrence of bacteria actively fixing carbon and nitrogen and thermophilic archaea in the rock habitat. Pacearchaeota has ecological advantages in colonizing the chimney rock interior, because the availability of nutrients and space is limited by silica deposition from hydrothermal fluids. We propose that the diversification of rock-hosted DPANN archaea could be profoundly influenced by coexisting microbes and minerals.

---

The archaeal superphylum DPANN (originally proposed by five phyla: Diapherotrites, Parvarchaeota, Aenigmarchaeota, Nanohaloarchaeota, and Nanoarchaeota) was first recognized by single-cell genomics^1^. Complete and near-complete DPANN genomes, including extra major groups such as Micraarcheota, Pacearcheota, and Woesearchaeta, are obtained by metagenomic assembly and binning into metagenome-assembled genomes (MAGs)^2–4^. Based on comparative genomic analysis, DPANN archaea have small genomes encoding a minimum of metabolic enzymes^2,5^. Thus, DPANN archaea are thought to depend on other microbes for most metabolites. By direct cell observations, along with phylogenomic profiling, DPANN archaea are attached to host archaea cells in fluid samples from co-cultures^6–9^, acid mine drainages^3, 10, 11^, and shallow and deep aquifers^12, 13^. However, recent studies have suggested that DPANN archaea may belong to complex microbial communities without symbiotic lifestyles^14, 15^.

Novel archaeal lineages referred to as deep-sea hydrothermal vent euryarchaeota (DHVE) were first recognized by 16S rRNA gene sequence analysis^16^. Genome-based taxonomic classification reveals that DHVE-3, DHVE-5, and DHVE-6 are affiliated with Aenigmarchaeota, Woesearchaeota, and Pacearchaeota, respectively^17^. In deep-sea hydrothermal fields, DPANN-affiliated MAGs are obtained from vent fluids^18^, sediments^19^, seafloor, and sub-seafloor metal sulfide deposits^20, 21^. It is challenging to clarify biological and environmental factors controlling the growth of DPANN in spatiotemporally fluctuated habitats. In a metal sulfide chimney without fluid venting (hereafter called an extinct chimney) from the Southern Mariana Trough, we discovered the dense colonization of ultra-small microbial cells, where DPANN dominance was revealed by 16S rRNA gene sequence analysis ^22^. To unveil the metabolic and ecological features of DPANN, we investigate microbial communities in the metal sulfide chimney by synchrotron-based X-ray spectroscopy, single-cell-level infrared spectroscopy, and genome-resolved metagenomic and metaproteomic analyses.

## High-cell density with Pacearchaeota dominance in the inner chimney wall

The extinct chimney was collected using the remotely operated vehicle Hyper-Dolphin at the Pika site^23^. The water depth and temperature of the sampling point were 2787 m and 1.7°C, respectively. The dense colonization of ultra-small cells was observed at the grain boundaries of chalcopyrite (CuFeS_2_), which makes up the inner chimney wall^22^. Microscopic observations of a 20-μm-thick section and subsequent image analysis revealed that the chalcopyrite grain boundaries have porosities of 15%–46% with an average pore throat radius of 2–5.5 μm in the inner chimney wall (Extended Data Fig. 1). Estimated permeabilities of 1 × 10^-15^–6 × 10^-14^ m^2^ fall within the range of sandstone^24^, which potentially renders O_2_ into the chimney wall from seawater through grain boundaries. However, this is not the case for the inner chimney wall, given our previous data showing that cuprite (Cu_2_O), a mineral readily dissolved in contact with oxygenated seawater, is formed in the grain boudaries^22^.

To clarify whether the chalcopyrite grain boundaries are not highly permeable because of mineral stuffing, synchrotron-based scanning fluorescence X-ray microscopy was used to visualize the distributions of elements such as C, N, O, Al, Si, Fe, and Cu with a soft X-ray beam in a diameter range of 0.6–0.8 μm (Fig. 1a). It was revealed that the chalcopyrite grain boundaries in the inner and central regions of the chimney wall are primarily filled with SiO_2_ (Figs. 1b and c and Extended Data Fig. 2). It was also revealed that the high densities of C and N are distributed along the SiO_2_-infilling grain boundaries of chalcopyrite. Thus, we confirmed that the chimney wall is less permeable than the grain boundaries of chalcopyrite without the infilling of silica and C- and N-bearing materials.

**Fig. 1.**
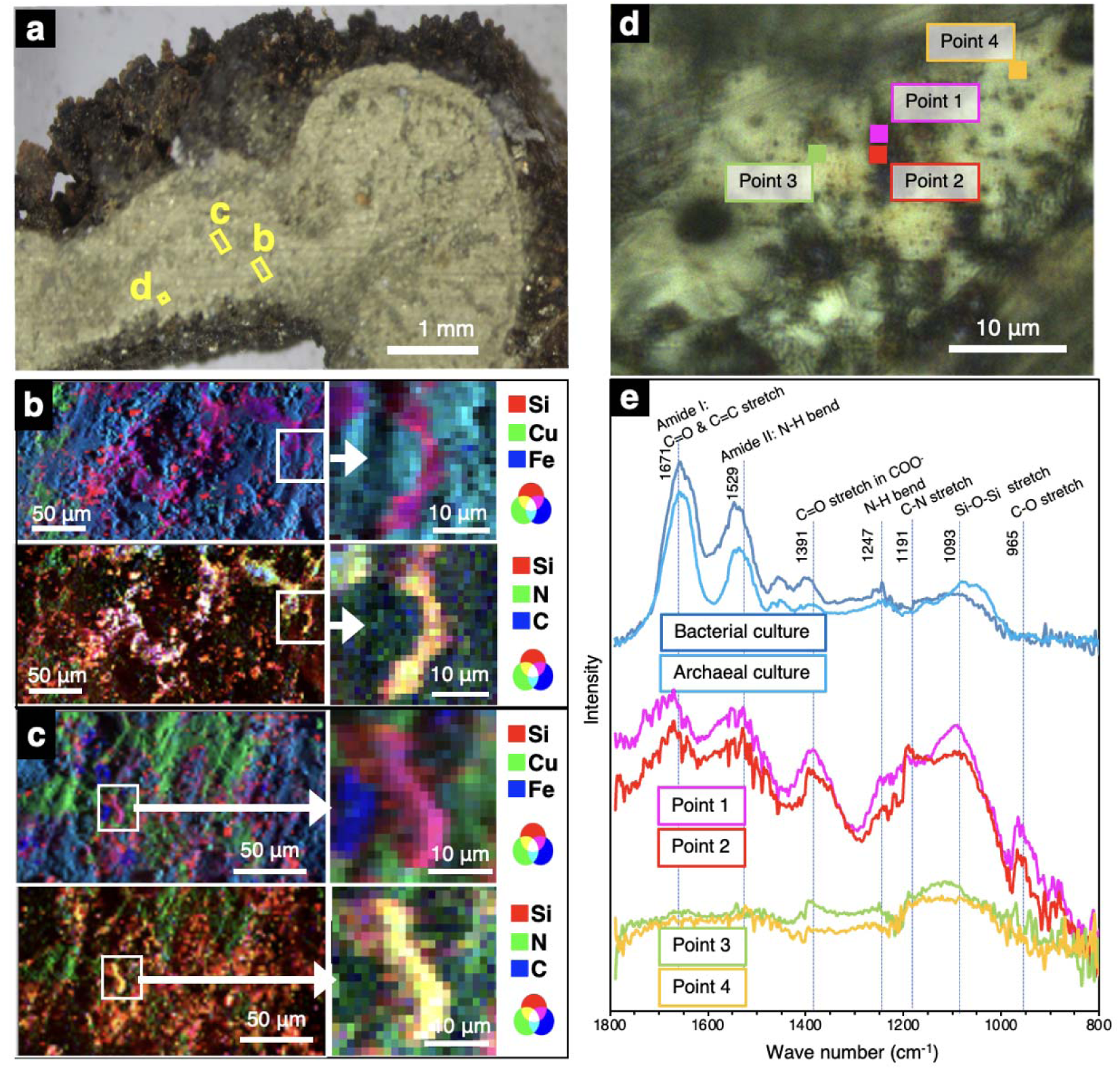
Submicron-scale spectroscopic analyses of chalcopyrite grain boundaries. **a,** A photograph of the inner chimney wall after cutting with a precision diamond wire saw. **b,c,** Elemental mapping of chalcopyrite grain boundaries obtained by synchrotron-based X-ray fluorescence analysis (XRF) from the regions with yellow rectangles in **a**. RGB diagrams are shown by synthesizing the three colors. **d,** A photograph of chalcopyrite grain boundaries with points analyzed by an infrared (IR) microscope. **e,** Optical photothermal infrared (O-PTIR) spectra of the points in **d** and cultured cells of *Nanobdella aerobiophila* strain MJ1^T^ (= JCM33616^T^) and *Metallosphaera sedula* strain MJ1HA (=JCM33617) for an archaeal reference and *Shewanella oneidensis* strain MR-1^T^ (=ATCC 700550^T^) for a bacterial reference. Peak assignment was based on refs^26–28^ and references therein.

Next, the coexistence of microbial cells and SiO_2_ within the chalcopyrite grain boundaries^25^ was investigated by infrared (IR) spectroscopy at a submicron resolution. IR spectra included peaks diagnostic of microbial cells (amide I and II^26^) and SiO_2_ (Si-O-Si stretch^27^) around a chalcopyrite grain (Figs. 1d and e). In addition, a peak attributed to carboxylic groups at ∼1400 cm^-1^ was prominent in the spectra around the chalcopyrite grain, indicating the presence of extracellular polymeric substances commonly found in biofilms^28^.

*In situ* spectroscopic analyses demonstrated the coexistence of microbial cells and SiO_2_ in the grain boundaries. As the limit of detection of C and N by synchrotron-based XRF analysis is ∼1000 ppm^29^, cell densities along the grain boundaries appear to be high. Consistently, the bulk cell density of the chimney interior primarily composed of the inner chimney wall (1.9×10^8^ cells/cm^3^) was as high as those in metal sulfide chimneys from the Central Indian Ridge and the western Pacific^30^. Before performing metagenomic analysis, two DNA extraction methods, respectively, based on physical (beads beating) and chemical (alkaline heating) rapturing methods were evaluated for the chimney interior sample. DPANN dominance was confirmed, regardless of the methods used (Extended Data Fig. 3 and Extended Data Table 1). Phylogenetic analysis of DPANN-affiliated 16S rRNA gene sequences classified the dominant DPANN as Woesearchaeota.

## Genomic features of the chimney Pacearchaeota

As the physically disrupted DNA extractant is less damaged than that chemically disrupted^30^, the former was subjected to genome-resolved metagenomic analysis^30^. From the chimney interior sample, near-complete genomes were reconstructed by several binning protocols (labeled as care and meta in genome names). By the phylogenetic analysis of 16 rRNA gene sequences in the reconstructed genomes and from the amplicon libraries, the reconstructed genomes named Idc_in_care_mg3 and Idc_in_meta_mg8 contained the 16S rRNA gene sequences nearly identical to the dominant DPANN lineage obtained by the amplicon sequence analysis (Extended Data Fig. 4 and Extended Data Table 2). The construction of a 122-concatenated-protein phylogenetic tree using the reconstructed genomes further confirmed the phylogenetic affiliation within Pacearchaeota rather than Woesearchaeota (Fig. 2)^31^. Based on 52 single-copy genes (Extended Data Tables 3)^32^, the sizes of the reconstructed genomes were 0.72 and 0.88 Mbp, whereas their completenesses were 84.6% and 92.3%, respectively (Extended Data Table 2).

**Fig. 2.**
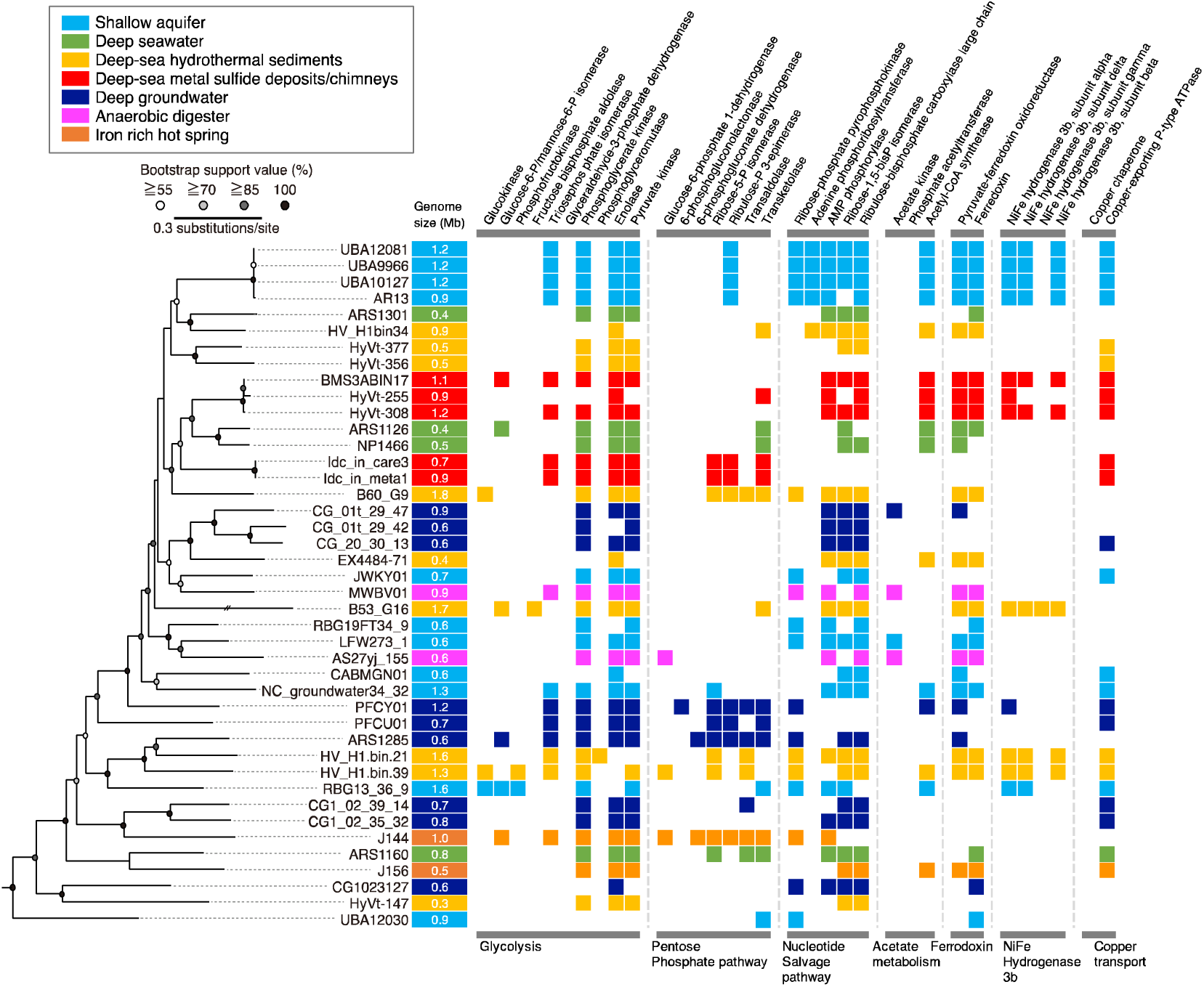
Comparative analysis of Paceacrhaeota-affiliated genomes from various habitats. On the left, a maximum likelihood phylogenetic tree of Pacearchaeota MAGs from this study (Idc_in_care_mg3 and Idc_in_meta_mg8) and from public databases was constructed using 122 archaeal marker genes with three Woesearchaeota MAGs as an outgroup (GW2011-AR15: GCA_000830295.1; GW2011-AR11:GCA_013330825.1; GW2011-AR9: GCA_013331635.1). On the right, an overview of the presence or absence of metabolic genes commonly found in Pacearchaeota MAGs is shown. Gene annotation was performed using BlastKOALA with default thresholds (see methods).

In the Pacearcaeota genomes, nearly full sets of genes involved in glycolysis and the non-oxidative pentose phosphate pathway (PPP) were encoded, indicating adenosine triphosphate (ATP) generation via fermentation of sugar-based biomolecules (Fig. 2 and Extended Data Table 4). Many genes involved in nucleotide biosynthesis, particularly pyrimidine bases, were present in the chimney Pacearchaeota genomes (Extended Data Table 5). In contrast, genes involved in the biosynthesis of amino acids, lipids, and cofactors were absent. Furthermore, metabolic capabilities other than carbon metabolism are lacking (Extended Data Table 6). The limited biosynthetic capabilities are linked to the symbiotic lifestyle of the chimney Pacearchaeota.

## Comparison of the Pacearchaeota genomes from various habitats

Although Pacearchaeota is ubiquitous across Earth’s ecosystems, its rarity and lack of obvious symbiotic partners make the lifestyle of Pacearchaeota enigmatic. As Pacearchaeota lineages across habitats and the genomic signatures of transitions among habitats remain unclear, we expand the inventory for Pacearchaeota genomes from natural and engineered environments such as anaerobic digesters^33, 34^, shallow and deep aquifers^12, 16, 32, 35, 36^, iron-rich hot spring^37^, deep seawater^38^, deep-sea hydrothermal sediments^21, 39, 40^, legacy radioactive waste trench water^40^, sub-seafloor metal sulfide deposits^20^, and deep-sea hydrothermal vent chimneys (Extended Data Table 2).

High-quality genomes with completeness higher than 60% were profiled with respect to biosynthetic and other metabolic capacities, as well as basic genomic features (Fig. 2 and Extended Data Tables 4–6). The 122-concatenated-protein phylogenetic tree of the Pacearchaeota genomes shows that the phylogenomic relationships were related neither to habitat type nor genome size, which was common in the 16S rRNA gene tree (Extended Data Fig. 4). The deep-sea metal sulfide deposits/chimneys hosted two distinct lineages with relatively high genome sizes (0.7–1.2 Mbp; Extended Data Table 2): one comprised the chimney genomes (Idc_in_care_mg3 and Idc_in_meta_mg8), and the other comprised genomes from metal sulfide deposits in the same deep-sea hydrothermal field (BMS3ABIN17, HyVt-255, and HyVt-308). The two lineages were clustered with deep-sea Pacearchaeota lineages with small genomes (ARS1126 and NP1466) with high bootstrap values. The lineage from the metal sulfide deposits appears to be metabolically flexible, based on the presence of genes involved in the nucleotide salvage pathway, Ni-Fe hydrogenase group 3b, and amino acid biosynthesis (Fig. 2 and Extended Data Tables 4–6). Muramoyltetrapeptide carboxypeptidase, which releases alanine from peptidoglycan^41^, a main component of the bacterial compartment, was found in abundance in the genomes of the chimney lineage (Extended Data Table 7).

## Genome annotations for the whole community in the chimney interior

To elucidate the biological interactions of Pacearchaeota with other co-occurring microbes, MAGs with completeness higher than 50% and contamination lower than 10% were profiled (Extended Data Tables 8 and 9). In addition to Pacearchaeota-affiliated genomes, we obtained nine taxonomically distinct genomes, five of which contained 16S rRNA gene sequences. As with Pacearchaeota, Patescibacteria (Idc_in_care_mg2 and Idc_in_meta_mg7) lacked biosynthetic and other metabolic capacities (Extended Data Tables 10–12). We found that Nitrosococcaceae (Gammaproteobacteria), the most abundant taxonomic group based on 16S rRNA gene amplicon sequences, was not found in MAGs (Extended Data Table 8). However, the Nitrosococcaceae sequences were expected to be taxonomically identical to gammaproteobacterial genomes classified as the order 21-64-14 without 16S rRNA gene sequences (Idc_in_meta_mg8 and 12). Hence, we constructed a phylogenetic tree based on 16S rRNA gene sequences from near-complete genomes of the order 21-64-14 in public databases and the Nitrosococcaceae-affiliated amplicon sequences (Extended Fig. 5 and Extended Data Table 13). As a result, the Nitrosococcaceae sequences were nearly identical to those from the order 21-64-14. Thus, the 21-64-14-affiliated genomes were confirmed to represent the abundant populations in the chimney interior.

## Metaproteomics resolved community-level metabolic activities

To understand the metabolic pathways and nutrient cycles potentially operated in the chimney interior, we annotated genes in the near-complete genomes from the dominant members of the whole community (Extended Data Tables 10-12). To constrain the partners of Pacearchaeota, it is crucial to clarify metabolically active members by metaproteomics. Duplicated extraction and sequencing led to 249 and 315 protein sequences, among which 133 and 219 protein sequences were obtained from the 21-64-14-affiliated genomes (Extended Data Tables 14 and 15). Relative abundances of the 21-64-14-affiliated genomes estimated by normalized genome coverages were similar to those estimated by 16S rRNA gene amplicon sequences and metaproteomic sequences (Fig. 3a). The 21-64-14-affiliated genomes included genes encoding a sulfate adenylyl transferase gene (sat), and anadenylyl-sulfate reductase genes (aprAB), and an array of dissimilatory sulfite reductase (dsr) genes. dsrC is involved in the reaction with dsrAB and sulfite and thought to be reduced by dsrMK(JOP)^42–44^. The presence of sat, aprAB, and dsr genes in the genome is known for dissimilatory sulfate reduction and sulfur oxidation. Sulfur oxidation is typically mediated by microbes, including dsrEFH for reactions with DsrC^45^ ^46^. The presence of dsrMK(JOP) and the absence of dsrEF support the inference that the 21-64-14-affiliated bacteria mediate dissimilatory sulfate reduction (Extended Data Tables 12). Metaproteomic analysis revealed that all genes involved in dissimilatory sulfate reduction, except for dsrM and dsrP, were detected in the chimney interior (Extended Data Table 16). In addition, metaproteomic detection of key enzymes for fixing carbon and nitrogen, such as Form I ribulose bisphosphate carboxylase/oxygenase (RuBisCO) large subunit, nitrogenase molybdenum-iron protein alpha chain (nifD), and nitrogenase iron protein (nifH) revealed the active primary production by the 21-64-14 bacteria (Fig. 3b and Extended Data Table 16).

**Fig. 3.**
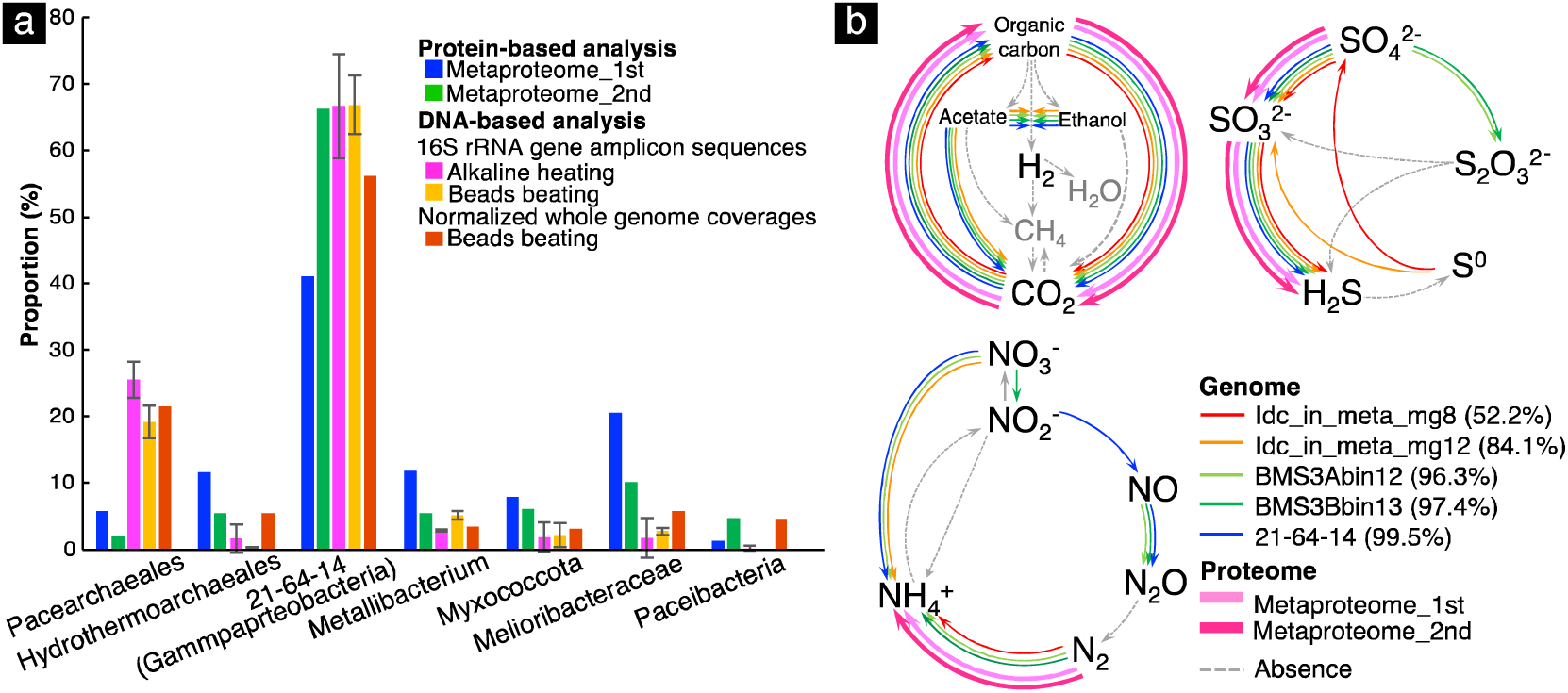
Taxonomic abundances and metabolic profiles of the chimney community. **a,** Comparison of taxonomic abundances based on protein- and DNA-based analyses. Proportions were obtained using selected data shown in Extended Data Tables 1 (16S rRNA gene amplicon sequences), 8 (normalized whole genome coverages), 14, and 15 (proteomics). **b,** Metabolic pathways/genes involved in biogeochemical cycles of C, N, and S found in the genomic and proteomic inventories of 21-64-14-affiliated bacteria. A percentage in parentheses indicates genome completeness.

In contrast to the major occurrence of Pacearchaeota in the 16S rRNA gene amplicon sequence libraries and normalized whole genome coverages, Pacearchaeota-affiliated sequences were minor in the metaproteomic libraries (Fig. 3a). This difference may be reasonable, considering the small genome sizes of Pacearchaeota. Pacearchaeota-affiliated protein sequences were annotated as hypothetical proteins, translation initiation factor IF-2 subunit alpha, and pyruvate kinase (Extended Data Tables 14 and 15). Pyruvate kinase catalyzes the last step of glycolysis to produce ATP and pyruvate from AMP, phosphoenolpyruvate, and phosphate for energy conservation^47^. The minor occurrence of Hydrothermoarchaeales in the 16S rRNA gene amplicon sequence libraries and normalized whole genome coverages was inconsistent with the frequent occurrence of Hydrothermoarchaeales-affiliated sequences in the duplicated metaproteomic libraries (Fig. 3a). Both proteomic libraries included Hydrothermoarchaeales-affiliated sequences annotated as archaeal chaperonin known to catalyze protein folding at high temperatures^48^ and enolase involved in the last second step of glycolysis^49^.

## Biological interactions of Pacearchaeota with potential partners

Pacearchaeota with the smallest median genome size across DPANN probably adopts an obligate symbiotic lifestyle, given their highly reduced biosynthetic and metabolic capacities^2, 5^. However, partner organisms remain unknown, primarily because of the minor occurrence of Pacearchaeota in taxonomically diverse communities. As many Pacearchaeota genomes include genes encoding murein transglycosylase (GH23 in the CAZy database) that specifically binds and degrades peptidoglycan, a symbiotic relationship with bacteria is speculated^50^. Although murein transglycosylase was absent in the chimney Pacearchaeota genomes (Extended Data Table 17), muramoyltetrapeptide carboxypeptidase, which releases alanine from peptidoglycan^41^, was found in abundance in the Pacearchaeota genomes from the chimney and various habitats (Extended Data Table 7). Metaproteomic evidence of bacterial primary production in the chimney interior agrees with the bacterial symbiosis with Pacearcheota. The successful enrichment of DPANN with only bacteria from deep-sea hydrothermal vent fluids also supports this possibility^18^.

The isoprenoid biosynthesis genes for archaeal mevalonate were present in the Hydrothermarchaeales genomes (Extended Data Table 10), indicating the supply of archaeal lipids from Hydrothermarchaeales to Pacearchaeota. Although all Hydrothermarchaeales genomes, including the chimney Hydrothermarchaeales, support chemolithoautotrophy via the Wood Ljungdahl pathway^51, 52^, cellular maintenance against heat stress rather than the active incorporation of inorganic carbon for the growth was clarified through our metaproteomic analysis.

## Ecological factors controlling the growth of Pacearchaeota

The sampling site is associated with the venting of black smokers (>300°C)^23^. The inner wall of chalcopyrite is universally formed in deep-sea hydrothermal vent chimneys, because the solubility of chalcopyrite decreases significantly at <300°C^53^. Although chalcopyrite grain boundaries are spatially available for microbes in the inner chimney wall, the temperature range of black smokers inhibits microbial colonization. After the transition from high- to low-temperature fluids, chemolithotrophic microbes appear to be hosted in the chalcopyrite grain boundaries. To distinguish microbial populations energetically depending on low-temperature fluids, the optimal growth temperature was estimated for the chimney community based on the frequencies of the seven amino acids Ile, Val, Tyr, Trp, Arg, Glu, and Leu (IVYWREL) in whole genome sequences^54^ (Fig. 4a and Extended Data Table 8). The optimal growth temperature of Hydorothermarchaeales was ∼70°C, consistent with the abundance of Hydorothermarchaeales in Juan de Fuca Ridge flank crustal fluids at 67°C^52^. Unexpectedly, the 21-64-14-affiliated genomes from the chimney sample had high frequencies of IVYWREL (estimated OPTs of ∼67°C), as well as some representative genomes of the 21-64-14 order (Extended Data Table 13). Thus, it is likely that the chemolithotrophy on low-temperature fluids is passed down from Hydorothermarchaeales to the 21-64-14 order. After the cessation of fluid venting, the cold chimney interior is limited to energy sources. The low OPTs of Pacearchaeota (∼44°C) and Patescibacteria (35°C) imply the nutritional dependence of the thermophilic primary production during fluid venting.

**Fig. 4.**
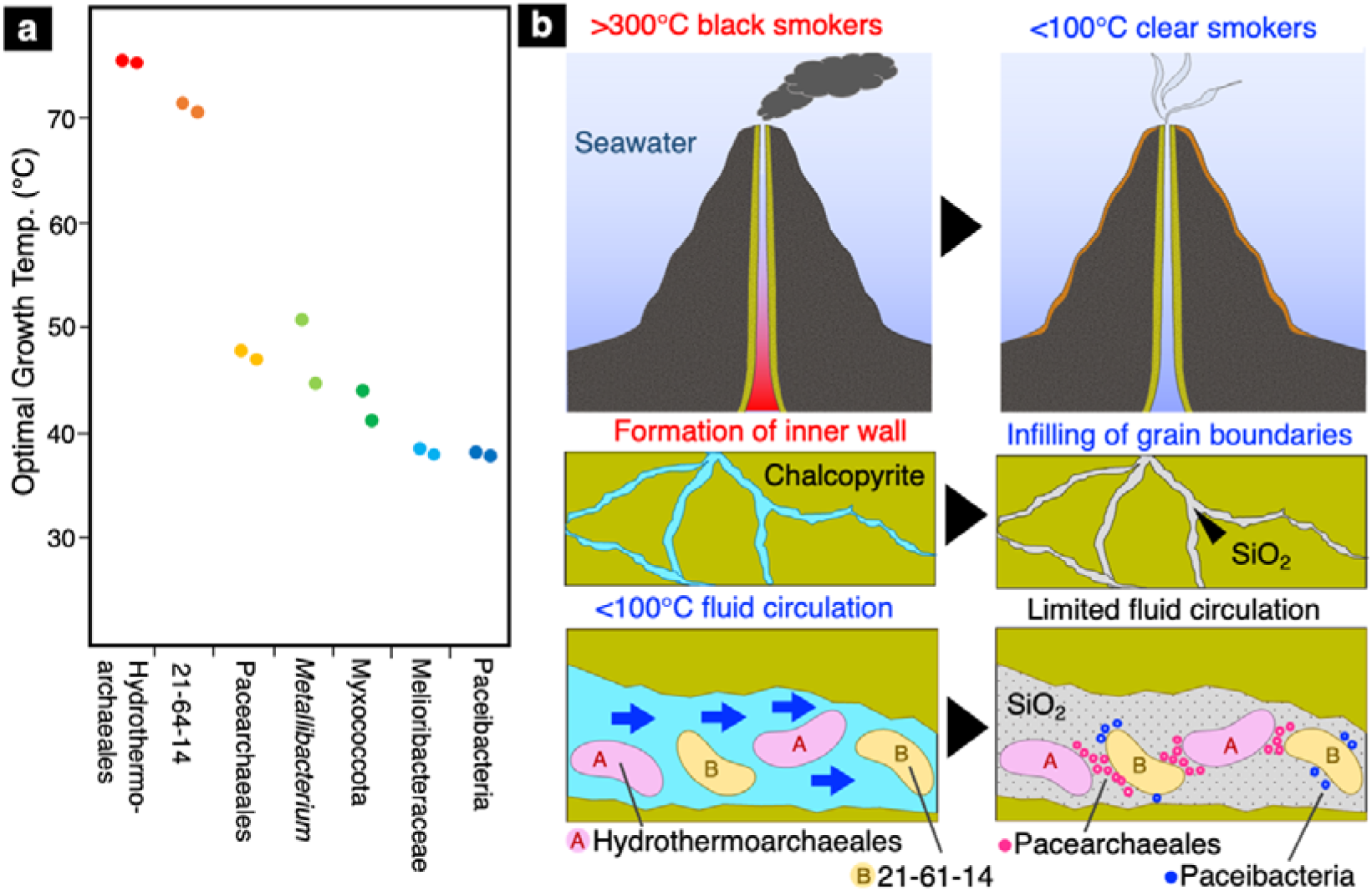
Temperature dependence of mineral formation and microbial colonization in a deep-sea hydrothermal vent chimney. **a,** Optimal growth temperatures of dominant members of the chimney community predicted by the frequencies of 7 amino acids (IVYWREL). **b,** Schematic illustrations of fluid venting, mineral formation, and key microbial populations. The blue arrows indicate the flow of low-temperature fluids.

As shown in Fig. 4b, amorphous silica is ubiquitously formed inside the chimney during the transition from high- to low-temperature fluid venting^55^. Thus, the colonization of the thermophilic populations is accompanied by the deposition of amorphous silica in chalcopyrite grain boundaries. As a result, pore size reduction favors the colonization of small cells. In the chalcopyrite grain boundaries, the chimney Pacearhaeota probably coexists with the assemblage of amorphous silica and thermophilic biomass (Fig. 4b). In our previous study, the occurrence of ultra-small cells (∼100 nm in size) was demonstrated in the chalcopyrite grain boundaries, with amorphous silica being mineralogically identified^22^. Glycolysis and the PPP, the central metabolism of the Chimney Pacearchaeota revealed by genome-resolved omics, are catalyzed by Fe(II) without enzymatic activities^56, 57^. Given the Fe(II) availability from the dissolution of chalcopyrite for non-enzymatic reactions, the chimney Pacearchaeota has some advantage in reducing protein production under energy-limited conditions.

## Evolutionary implications

We revealed the rock-hosted lifestyle of Pacearchaeota influenced by the microbe-mineral assemblages in the deep-sea hydrothermal vent chimney. Sinters are primarily composed of amorphous silica universally formed at terrestrial hot springs in association with archaeal colonization^58^. Hydrothermarchaeota, formally known as Marine Benthic Group E, is also widespread in terrestrial hot springs^52, 59^. Hydrothermarchaeota is an early-evolved archaeal phylum branching between the DPANN and Euryarchaeota groups^51^. Based on the 27 highest-ranked marker genes, DPANN is also found in a deeply branched archaeal group on the new tree of life^60, 61^. The relationship of deeply branching archaeal lineages demonstrated in this study could represent an evolutionary model that drives the archaeal diversification at hydrothermal vents.

## Acknowledgements

We thank captain and crews of the R/V Natsushima and the ROV Hyper-Dolphin operating groups for their technical support in sample collection. We also thank to the scientists that joined the NT12-24 cruise and to the members of TAIGA project for providing opportunity of this study. We are grateful to Hitoshi Furutani, Koji Ichimura, Hanae Kobayashi, Makoto Momose and Miho Hirai for their technical assistance. We thank Edanz Group (https://en-author-services.edanz.com/ac) for editing a draft of this manuscript.

## Author contributions

H.T., M.K., and Y.S. designed the study. Y.S. and S.K. collected and analyzed the chimney sample as shipboard scientists during JAMSTEC Scientific Cruises NT12-24. T.Y. performed image analysis for porosity and permeability. H.S., M.O., M.K., and Y.S. performed synchrotron-based spectroscopy. Y.S. performed IR spectroscopy. H.T. M.K., S.K., and Y.S. performed single-gene and metagenomics analyses. H.T. M.K., M.S., A.K., M.M, and Y.S. performed metaproteomic analysis. H.T., M.K., and Y.S. co-wrote the manuscript. All authors discussed the results and commented on the manuscript. The synchrotron radiation experiments were conducted at RIKEN Coherent Soft X-ray Spectroscopy Beamline (BL17SU) in SPring-8 with the approval of JASRI (Proposal No. 2022B1531).

## Competing interests

The authors declare that they have no competing interests.

## Additional information

**Extended data and Supplementary Tables** are available for this paper at https://doi.org/XXXX/XXXX-XXX-XXX-X.

**Correspondence and requests for materials** should be addressed to Y.S.

## Data availability

All data required to evaluate the conclusions in the paper are present in the paper and/or the Supplementary Materials. The 16S rRNA gene sequences in this study were all deposited in the DDBJ nucleotide sequence database with accession numbers XXXXXXXX-XXXXXXXX. The MAG sequences were deposited under the BioProject accession number PRJDB13464.

## Methods

### Sample collection and subsampling

From an active hydrothermal vent field at the Pika site (12°55.15’N, 143°36.96’E) in the Southern Mariana Trough, a metal sulfide chimney sample was collected during the Japan Agency for Marine-Earth Science and Technology (JAMSTEC) Scientific Cruise NT12-24 of the R/V Natsushima (September in 2012). The metal sulfide chimney sample was placed in a container from the surrounding seawater during transportation to the surface using the manipulator arm of the remotely operated vehicle (ROV) Hyper-Dolphin. It was visually confirmed that the chimney structure lacked hydrothermal fluid venting. For the subsampling of the interior portion, the chimney exterior was removed using sterile chisels and spatulas after flaming with a gas torch to prevent contamination from the exterior potion. An intact portion of the chimney interior was preserved with 3.7% formamide in seawater onboard and then substituted with 50% ethanol in distilled, deionized water for spectroscopic characterizations. The interior portion of the sample ground into powder by sterile pestles and mortars was frozen at -80°C for total cell counting and metagenomic and metaproteomic analyses.

### Image analysis of chalcopyrite grain boundaries for the estimation of permeability

A 20-μm-thick section previously characterized^22^ was subjected to image analysis. The permeability *k* (m^2^) was predicted with good accuracy using the following equation^62^, *k* = 8.5(*ϕr*^2^)^1.3^, where *ϕ* (dimensionless) denotes the fraction of pores open to the outside, and *r* (m) represents the radius of the narrowest portion of the pore penetrating through the sample (pore throat). Image analysis was performed using ImageJ^63^. Each of the two analyzed areas in the Extended Data Fig. 1 was first converted to grayscale and then binarized (white: chalcopyrite grain; black: grain boundary). When using the algorithms provided with ImageJ for binarization thresholding, *ϕ* was overestimated in many cases (*ϕ* = 0.46-0.60). Alternatively, the minimum value *ϕ* was estimated to be 0.15 when determined visually. Therefore, the range of *ϕ* was set to be 0.15–0.46. Although *r* represents the pore throat, it is difficult to determine this value accurately in a two-dimensional image. Therefore, after extracting a relatively long connected path using ImageJ, *r* was visually estimated by assuming that the typical radii of the path (2–5.5 µm) can approximate *r*. Substituting *r* = 2 μm and *ϕ* = 0.15 for the minimum value of *k* and *r* = 5.5 μm and *ϕ* = 0.46 for the maximum value of *k* into the above equation yields *k* = 1 × 10^-15^–6 × 10^-14^ m^2^.

### Spectroscopic characterizations of the chimney interior

To clarify the distributions of microbes and minerals, the intact portion of the chimney structure was cut into 3-mm-thick sections using a precision diamond wire saw (Meiwa Fosis Corporation DWS 3500P). As the surface of the thin section was smooth, no further polishing was required for subsequent spectroscopic characterizations.

Soft X-ray spectromicroscopy developed at the soft X-ray undulator beamline BL17SU of SPring-8 was used for mapping light to heavy elements (C, N, O, Al, Si, Fe, and Cu) at a submicron resolution^29^. The incident soft X-ray beam was focused using the Fresnel zone plate. For micro XRF, the energy and intensity of the fluorescent soft X-rays emitted from the sample were analyzed using a silicon drift detector. For simultaneous measurements of seven elements (C: 120–170 ch; N: 185–240 ch; O: 268–313 ch; Fe: 368–410 ch; Cu: 480–540 ch; Al: 767–843 ch; Si: 920–960 ch), an excitation energy of 2100 eV was used. To obtain XRF spectra to resolve a peak of N from that of O, the excitation energy was lowered to 475 eV.

A mIRage infrared microscope (Photothermal Spectroscopy Corp., Santa Barbara, USA) was used to acquire O-PTIR spectra at a submicron resolution in refection mode (Cassegrain 40 objective (0.78 NA)) with a continuous wave (CW) 532-nm laser as the probe beam. The pump beam consisting of a tunable QCL device (950–1800 cm^-1^; 2-cm^-1^ spectral resolution and 10 scans per spectrum) or a tunable OPO device (2700–3592 cm^-1^; 4-cm^-1^ spectral resolution and 1 scan per spectrum) was used to obtain O-PTIR spectra over the mid-IR ranges. Co-cultured cells of Nanoarchaeota strain MJ1 and *Metallosphaera* sp. strain MJ1HA (JCM33617) and cultured cells of *Shewanella oneidensis* (ATCC 700550) were lyophilized, and O-PTIR spectra were obtained by mounting on disks made of CaF_2_.

### Total cell count

The total number of cells in the chimney interior was measured using a direct count method with SYBR Green I. The formalin-fixed powdered sample was suspended in a 1:1 ethanol/phosphate-buffered saline solution. Some portions of the suspensions were sonicated for 30 s. A 0.22-µm-pore-size, 25-mm-diameter polycarbonate filter (Millipore) was used to collect the sonicated suspension. To stain the microbial cells, the filter was incubated in a TAE buffer containing SYBR Green I for 5 min at room temperature. The stained filter was briefly rinsed with deionized water and observed under epifluorescence using the Olympus BX51 microscope with the Olympus DP71 CCD camera.

### DNA extraction and 16S rRNA gene amplicon analysis

Prokaryotic DNA was extracted from the frozen powdered chimney subsample using chimerical and physical disruption methods. For chemical disruption^64^, the powdered chimney subsample was incubated at 65°C for 30 min in an alkaline solution consisting of 0.5-M NaOH and TE buffer (Nippon Gene Co.). After centrifugation at 5,000×g for 30 s, the supernatant was neutralized with 1-M Tris–HCl (pH 6.5; Nippon Gene Co.). After neutralization, the DNA-bearing solution (pH 7.0–7.5 in TE buffer) was stored at −4°C or −20°C for longer storage. For physical disruption, a DNeasy PowerSoil Pro Kit (Qiagen) was used, according to the manufacturer’s instructions. For the negative control, DNA extraction was performed without the addition of the subsample.

The 16S rRNA gene sequences were amplified by PCR using LA Taq polymerase (TaKaRa-Bio, Inc., Shiga, Japan) using the primers Uni530F and Uni907R^65^. The PCR was performed in a reaction mixture containing 0.1-μM oligonucleotide primer and ca. 0.1-ng/μL DNA template with 35 cycles of denaturation at 96°C for 20 s, annealing at 56°C for 45 s, and extension at 72°C for 120 s. The first PCR product was used for the second PCR step, which was run for 10 cycles with Illumina TruSeq P5 and Index-containing P7 adapters, which was purified using a MinElute Gel Extraction Kit (Qiagen, Inv., Valencia, CA). 16S rRNA gene sequencing was performed using the Illumina MiSeq Reagent Kit v2 by Illumina MiSeq System (Illumina, San Diego, US). The paired-end sequence reads were demultiplexed, trimmed, quality filtered, and chimera removed using standard Illumina software. The screened reads were imported into Qiime2 version 2022.2^66^ and DADA2 version 1.5.2^67^. For alignment and taxonomic affiliation of the representative sequence obtained, SILVA SSU Ref NR database version 138^68^ was used in the Qiime2 program.

### Genome-resolved metagenomic analysis

We used genomic DNA extracted by physical disruption for shotgun library construction with a KAPA Hyper Prep kit for Illumina (KAPA Biosystems)69. The sequencing of the library was performed on an Illumina MiSeq platform (MiSeq PE300). To reconstruct MAGs, the Read_QC module included in MetaWRAP v.1.3.2^70^ was used to trim and filter reads from the library. SPAdes version 3.13.0 with the options “—meta, -k 55,77,99,111,121”^71^ was used to assemble high-quality reads into contigs, which were binned into MAGs using the Binning module (including metabat2 MAGs using the Binning module (including metabat2^72^, maxbin2^73^ and concoct^74^) included in MetaWRAP. The reconstructed MAGs were further refined using the Bin refinement module in MetaWRAP. The gene prediction and annotation”of t’e)contigs and the near-complete genome were initially annotated using Prokka^75^. To estimate the relative abundances of MAGs, normalized read coverage values were calculated using the Quant bins module in MetaWRAP.

For phylogenomic analysis, we selected MAGs with completeness of >50% and contamination of <10% (Extended Data Table 8). The taxonomic classification was performed according to the Genome Taxonomy Database (GTDB) taxonomy^76^. 16S rRNA gene sequences in MAGs were taxonomically classified according to SILVA ver. 138^68^. Based on 120 concatenated single-copy marker proteins, a maximum likelihood tree was constructed for Pacearchaeota genomes. The GTDB database was used to align and trim the concatenated protein sequences^77^. The resulting alignment was trimmed using trim-al v1.4.1^78^ with the “-gappyout” option. The maximum likelihood phylogenetic tree was generated using W-IQ-TREE^79^ with 1,000 replicates of ultrafast bootstrap^80^ and visualized using Figtree v1.4.4 (http://tree.bio.ed.ac.uk/software/figtree/).

The Kyoto Encyclopedia of Genes and Genomes (KEGG) pathway tool^81^, along with the BlastKOALA tool^82^, was used for functional protein characterizations. We also used the KEGG Decoder to annotate protein sequences^83^. Curated sets of genes involved in hydrogen, carbon, nitrogen, and sulfur metabolism were searched using METABOLIC v.4.0^84^. In addition to KEGG, METABOLIC v.4.0 annotates genes using TIGRfam^85^, Pfam^86^, and custom hidden Markov model profiles.

### Metaproteomic analysis

The powdered chimney subsample was incubated in 12-mM SDC/12-mM SLS/100-mM TEAB buffer (pH 8.5) containing a 1% protease inhibitor (Protease Inhibitor Cocktail for General Use, Nacalai Tesque, Inc., Kyoto, Japan). After three cycles of sonication (30-s intervals, 0.5-s pulse on/0.5-s pulse off), the aliquot was cooled on ice. After centrifugation at 13,000×g for 3 min, the extracted proteins in lysis buffer (12-mM sodium deoxycholate, 12-mM sodium N-dodecanoylsarcosinate, and 50-mM ammonium bicarbonate containing 1% protease inhibitor cocktail for general use [Nacalai Tesque, Kyoto, Japan]) were reduced using 10-mM dithiothreitol at 37°C for 30 min following alkylation using 50-mM iodoacetamide at 37°C for 30 min in the dark. After five-fold dilution with 50-mM ammonium bicarbonate, the sample was digested using Lys-C (Wako, Osaka, Japan) at 37°C for 3 h following trypsin (Promega, Madison, WI, USA) at 37°C for 16 h. The digest acidified using trifluoroacetic acid was centrifugated at 15,000 × g for 1 min, the supernatant of which was desalted with C18-StageTips^87^ and dried under reduced pressure.

An ultra-high-performance nano-flow chromatography system and a hybrid trapped ion mobility spectrometry–quadrupole time of flight mass spectrometry system equipped with a nanoElute and a timsTOFPro (Bruker Daltonics, Bremen, Germany) were used for proteomic analysis. The dried sample was dissolved in formic acid (FA)/acetonitrile (ACN)/water (0.1/2/98, v/v/v) and then injected into a self-packed column (ACQUITY UPLC BEH C18, 1.7 µm, ID = 75 µm, length = 250 mm). For peptide separation, (A) FA/water (0.1/100, v/v) and (B) FA/ACN (0.1/100, v/v) were used at 60C as the mobile phase. The composition of the mobile phase (B) was changed at 2%–35% for 100 min, 35%–80% for 10 min, and 80%–80% for 10 min, maintaining a flow rate of 280 nL/min. The eluted peptides were analyzed using a parallel accumulation serial fragmentation scan mode^88^.

FragPipe (version 17.1) software was used to analyze liquid chromatography-mass spectrometry raw data^89^. Protein sequences of the MAGs reconstructed in this study (Extended Data Table 8) and 43 protein sequences considered contaminants (<u>cRAP protein sequences (thegpm.org)</u>) were used as reference databases. The enzyme was set to trypsin as a specific cleavage, and up to two missed cleavages were allowed in the proteolysis process. The allowed peptide lengths and mass ranges were 7–50 residues and 500–5,000 Da, respectively. Carbamidomethylation at cysteine residues was set as a fixed modification, whereas N-acetylation at the protein N-terminus and oxidation at methionine residues were set as variable modifications, allowing for up to three sites per peptide. The peptide spectrum matches and the identified peptides/proteins were determined at <1% false discovery rage (FDR) at the protein level.

---

**Extended Data Fig. 1.**
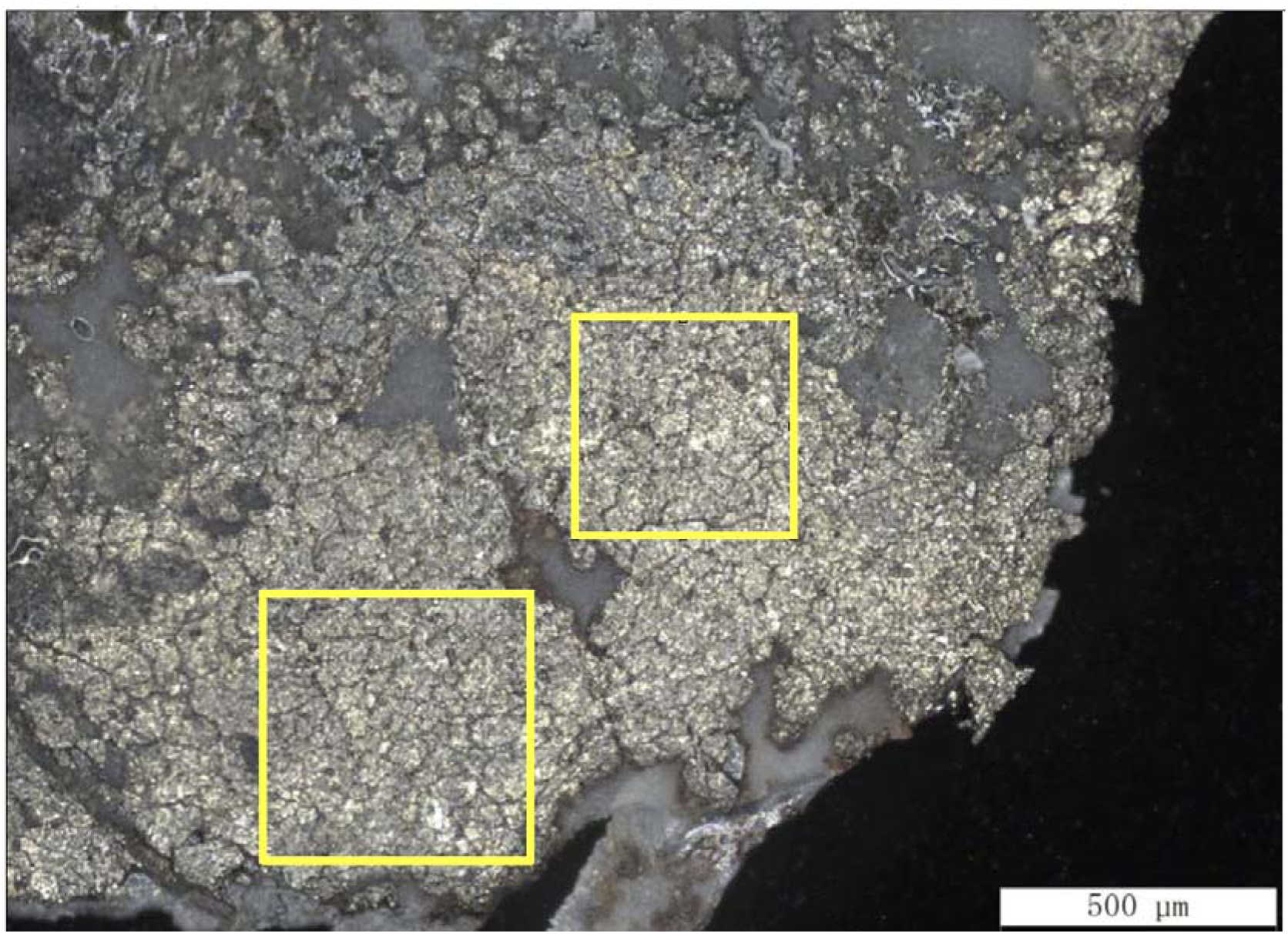
Chalcopyrite grain boundaries analyzed for porosity and permeability. A microscope image of a 20-μm-thick section from the inner chimney wall previously characterized^23^. The images in the yellow square were processed to measure porosity around chalcopyrite grains.

**Extended Data Fig. 2.**
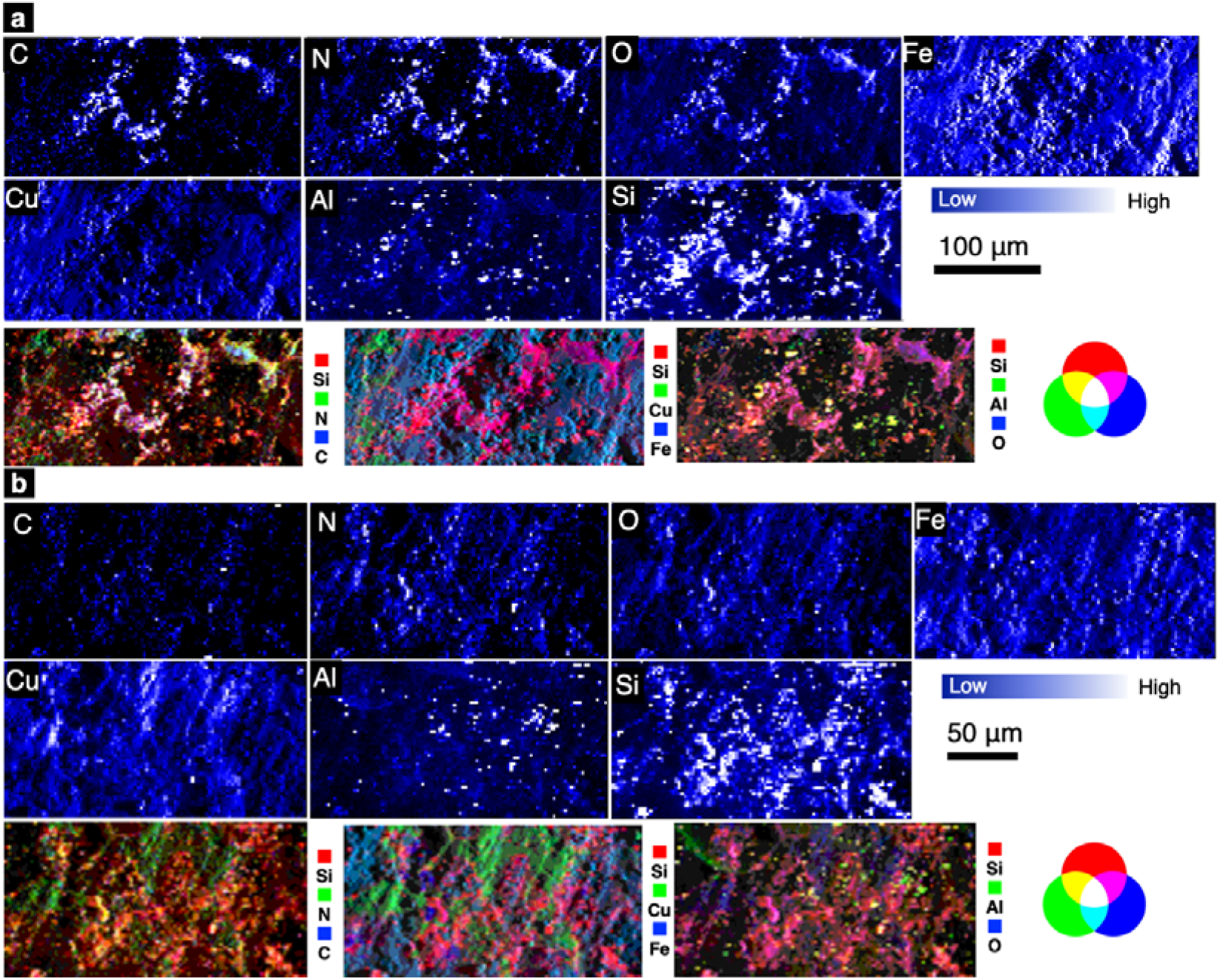
Elemental distributions of chalcopyrite grain boundaries. **a,b,** elemental maps obtained by synchrotron-based X-ray fluorescence analysis (XRF) with RGB color synthesis of selected elements from the regions with yellow rectangles labeled, respectively, with c and d in Fig 1a.

**Extended Data Fig. 3.**
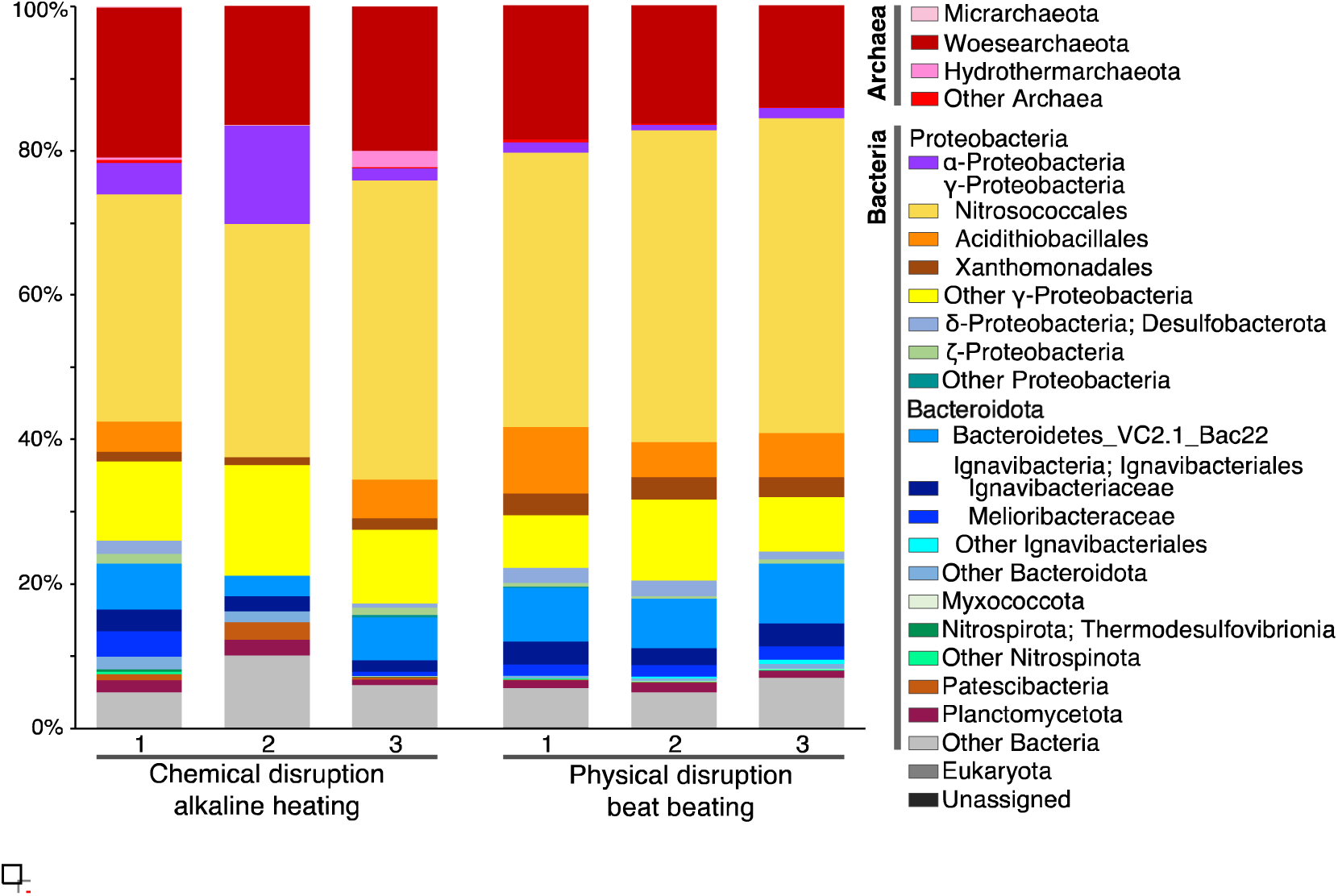
Microbial community structures from the chimney interior. Chemically and physically disrupted DNA extractants were analyzed for 16S rRNA gene amplicon sequences. The phylum and class were classified by prokaryotic 16S rRNA gene sequences with the nomenclature based on the SILVA 138 database using QIIME2 software.

**Extended Data Fig. 4.**
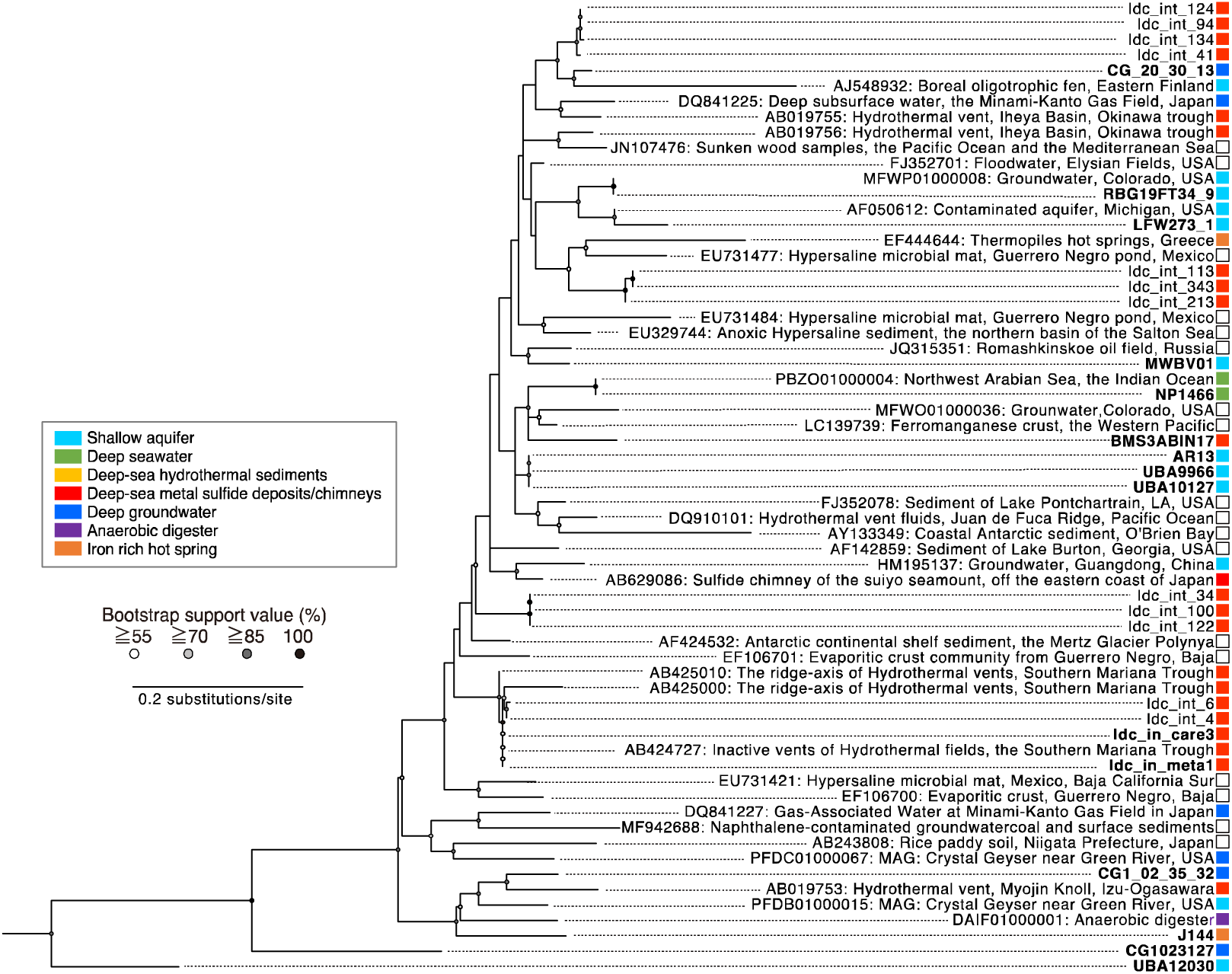
Analysis of Pacearchaeota-affiliated 16S rRNA gene sequences from the chimney interior and public databases. A maximum likelihood tree consists of 16S rRNA gene sequences from polymerase chain reaction (PCR) amplification with primers and from MAGs. 16S rRNA gene amplicon sequences obtained from the chimney interior in this study are prefixed with “Idc_int_,” whereas 16S rRNA gene sequences in MAGs are shown in bold text.

**Extended Data Fig. 5.**
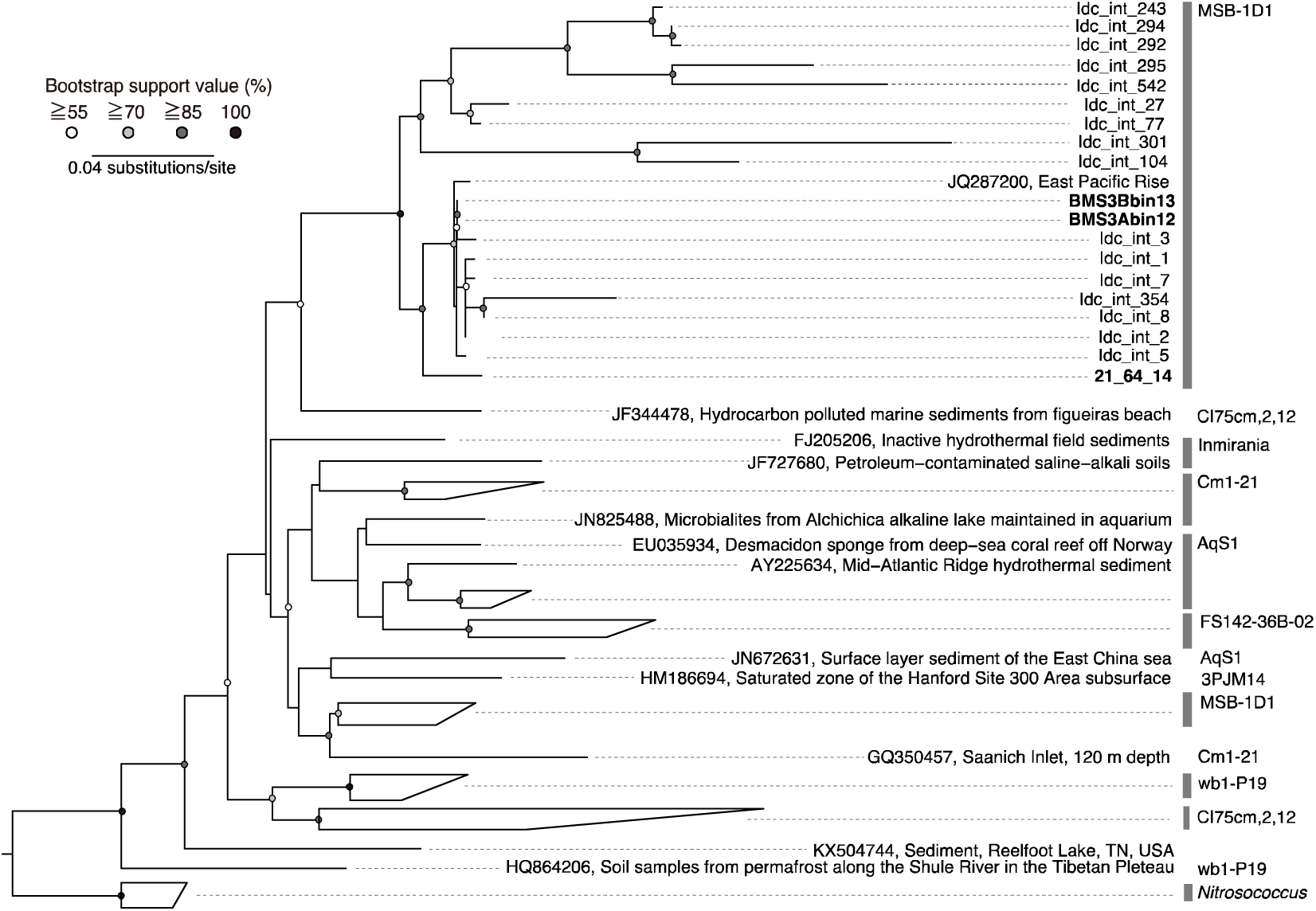
16S rRNA gene sequence analysis of Nitrosococcaceae- and 21-64-14-affiliated sequences. A maximum likelihood tree consists of 16S rRNA gene sequences from PCR amplification with primers and from MAGs. 16S rRNA gene amplicon sequences obtained from the chimney interior in this study are prefixed with “Idc_int_,” whereas 16S rRNA gene sequences in MAGs are shown in bold text.

## Notes

### Competing Interest Statement

The authors have declared no competing interest.

